# Mechanical stretch disrupts intracellular structures under impaired actin integrity in vascular smooth muscle cells

**DOI:** 10.64898/2026.05.17.725699

**Authors:** Eiji Matsumoto, Shinji Deguchi

**Affiliations:** Graduate School of Engineering Science, The University of Osaka, 1-3 Machikane-yama, Toyonaka, 560-8531, Osaka, Japan; Global Center for Medical Engineering and Informatics, The University of Osaka, Osaka, Japan; R^3^ Institute for Newly-Emerging Science Design, The University of Osaka, Osaka, Japan

**Keywords:** vascular smooth muscle cells, actin polymerization, Latrunculin A, mechanical stretch, actin-bundle organization, stochastic modeling

## Abstract

Actin-bundle organization is essential for vascular smooth muscle cell mechanics and is implicated in actin-related diseases. However, it remains unclear how cell stretching affects intracellular actin bundles when actin polymerization is impaired. Here, we performed live imaging of Latrunculin A-treated A7r5 vascular smooth muscle cells in a stretch chamber. GFP–***α***-actinin imaging showed that Latrunculin A reduced actin-bundle coverage while periodicity was maintained. Subsequent mechanical stretch disrupted both actin-bundle coverage and periodicity. We constructed a stochastic filament bundle model in which actin filament length, actin crosslinking protein dynamics, external stretch, and myosin-driven contractile shortening determine bundle connectivity. The model generated non-spanning, collapse, and persistent states based on spanning connectivity before and after stretch, shaped by filament length and applied strain. A reduced model further showed that these states are governed by a balance between connectivity formation and stretch-induced loss. Together, our results suggest that reduced actin polymerization destabilizes intracellular actin-bundle organization under mechanical stretch, providing a mechanism linking actin polymerization defects to mechanical fragility in vascular smooth muscle cells.

## 1 Introduction

Vascular smooth muscle cells are exposed to tensile forces in the vessel wall, where actin-based cytoskeletal structures contribute to bearing mechanical loads [1–3]. In skeletal muscle, excessive strain can cause muscle injury, in which muscle fibers and the surrounding extracellular matrix are partially disrupted before subsequent repair [4, 5]. By contrast, vascular smooth muscle is not typically described as undergoing an acute skeletal-muscle-strain-like rupture at the tissue level. Instead, mechanical fragility in these cells may emerge at the intracellular level through destabilization of force-bearing cytoskeletal organization [6–9]. This view is relevant to smooth muscle actinopathy, particularly diseases associated with mutations in ACTA2, which encodes smooth muscle *α*-actin [10, 11]. ACTA2 mutations have been identified as a major genetic cause of heritable thoracic aortic aneurysm and dissection, indicating that abnormalities in smooth muscle actin can impair vascular smooth muscle cell function [12, 13]. Mechanistic studies have shown that disease-associated ACTA2 mutations impair actin filament assembly, increase the fraction of unpolymerized actin, and reduce filament stability [14–16]. These findings raise the possibility that impaired actin polymerization may create a form of intracellular mechanical fragility in smooth muscle cells, even when the cell body itself does not undergo apparent rupture. Clarifying how actin polymerization defects alter intracellular force-bearing cytoskeletal organization is therefore essential for understanding cellular-scale mechanical fragility in actin-related vascular disease.

To examine the role of actin polymerization in smooth muscle cell mechanics under load, pharmacological perturbation provides a useful approach. Latrunculin A is a marine natural product originally isolated from sponges of the genus Latrunculia [17] and has been widely used in cell biology as an inhibitor of actin polymerization [18, 19]. It binds actin monomers stoichiometrically and prevents their incorporation into filaments [18]. Among latrunculin analogs, latrunculin B acts through a similar mechanism but is generally considered less potent than latrunculin A [20]. Although disease-associated ACTA2 mutations would provide a genetic model of smooth muscle actinopathy, introducing such mutations can induce broad changes in cytoskeletal organization, cellular phenotype, actin-isoform expression, and gene-regulatory state [15, 21, 22], making it difficult to isolate the role of reduced actin polymerization itself. By contrast, latrunculin A enables targeted weakening of actin filament formation without genetic modification. While latrunculin A has been widely used to perturb actin organization in vascular smooth muscle cells, its effects have often been examined under static or non-stretched culture conditions [23–25]. Thus, how cells with weakened actin filament formation respond to tensile loading remains unclear.

Here, we use Latrunculin A-treated A7r5 vascular smooth muscle cells to examine how reduced actin polymerization alters intracellular actin-bundle organization under mechanical stretch. We combine live-cell imaging of GFP–*α*-actinin in a stretch chamber with a stochastic filament bundle model and its reduced model to analyze a mechanism underlying stretch-dependent actin-bundle destabilization.

## 2 Results

### 2.1 Latrunculin A sensitizes actin bundles to mechanical stretch

We first examined GFP–*α*-actinin dynamics in A7r5 cells by live imaging in a stretch chamber (Fig. 1A). In a representative time-lapse experiment, GFP–*α*-actinin-positive bundle structures were partially reduced after Latrunculin A treatment and became fragmented after stretch, followed by partial reorganization after washout (Fig. 1B). We measured detected fiber area and periodicity using tile-based image analysis (Fig. 1C). Detected fiber area quantified GFP–*α*-actinin-positive bundle area, whereas periodicity reflected spatial ordering. Both metrics decreased when mechanical stretch was applied after Latrunculin A treatment.

**Fig. 1.**
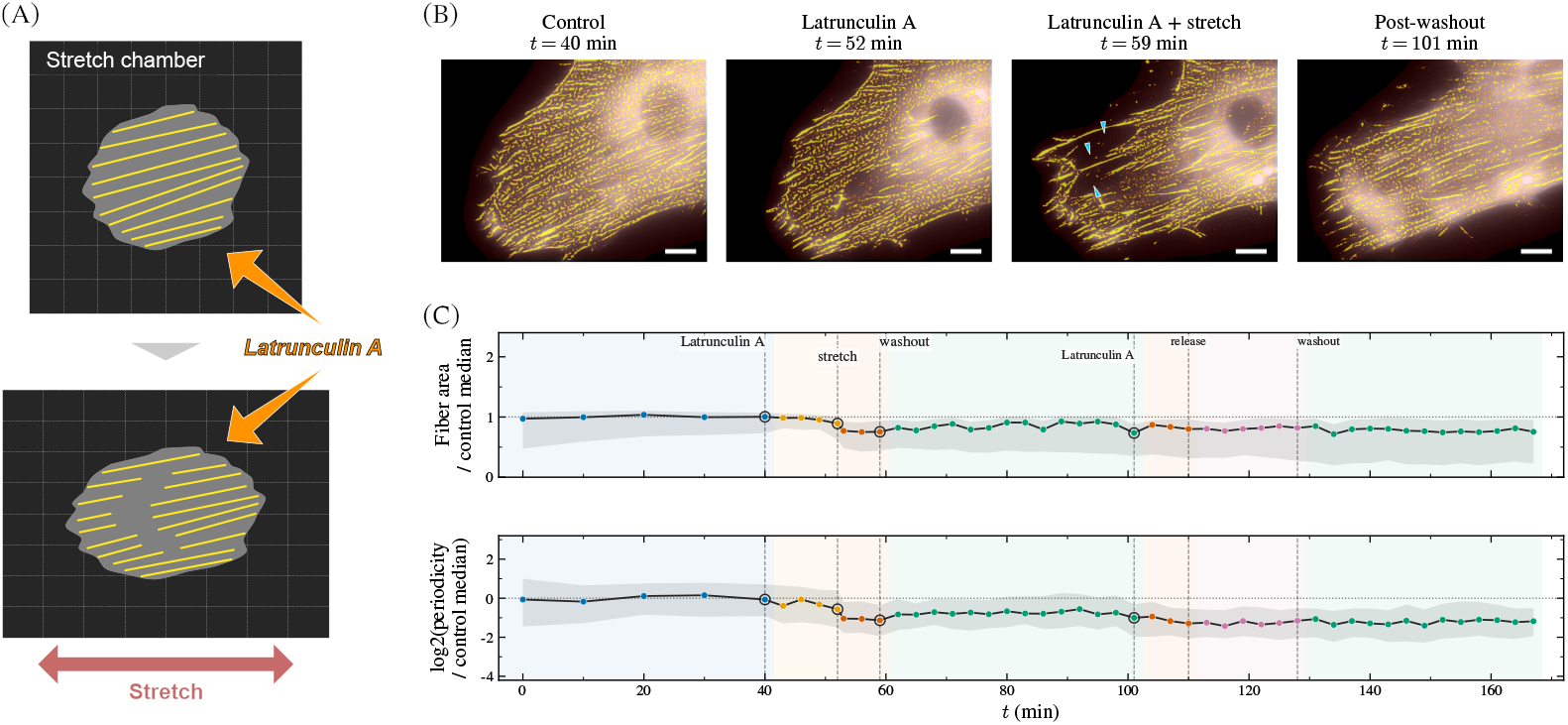
Latrunculin A treatment and mechanical stretch destabilize GFP– *α*-actinin-labeled actin-bundle organization. **(A)** Schematic of live-cell imaging using a stretch chamber during Latrunculin A treatment and mechanical stretch. **(B)** Representative GFP–*α*-actinin images from a single time-lapse experiment under Control, Latrunculin A, Latrunculin A + stretch, and post-washout conditions; the full time-lapse sequence is shown in Supplementary Video S1. Yellow overlays indicate detected GFP–*α*-actinin-positive bundle regions. Arrowheads indicate regions with reduced GFP–*α*-actinin-positive bundle continuity after stretch. Scale bars, 10 *µ*m. **(C)** Time courses of detected fiber area and periodicity. Metrics were normalized to the median of pre-Latrunculin A control tiles; periodicity is shown as a log_2_ ratio. Lines show tile medians, and shaded regions show interquartile ranges. Vertical dashed lines mark Latrunculin A addition, stretch, washout, release, and repeated stimulation events.

To quantify local bundle discontinuity, we performed manual line-ROI analysis along visible GFP–*α*-actinin-positive bundle-like structures (Fig. 2A and B). The maximum peak-to-peak gap was largest when stretch was applied after Latrunculin A treatment, whereas control, stretch-only, and Latrunculin A-only conditions showed comparable gaps (Fig. 2A). The corresponding collapsed ROI fraction showed the same trend (Fig. 2B). These results indicate that stretch after Latrunculin A treatment promotes local actin-bundle collapse, consistent with intracellular mechanical fragility of polymerization-weakened bundle organization.

**Fig. 2.**
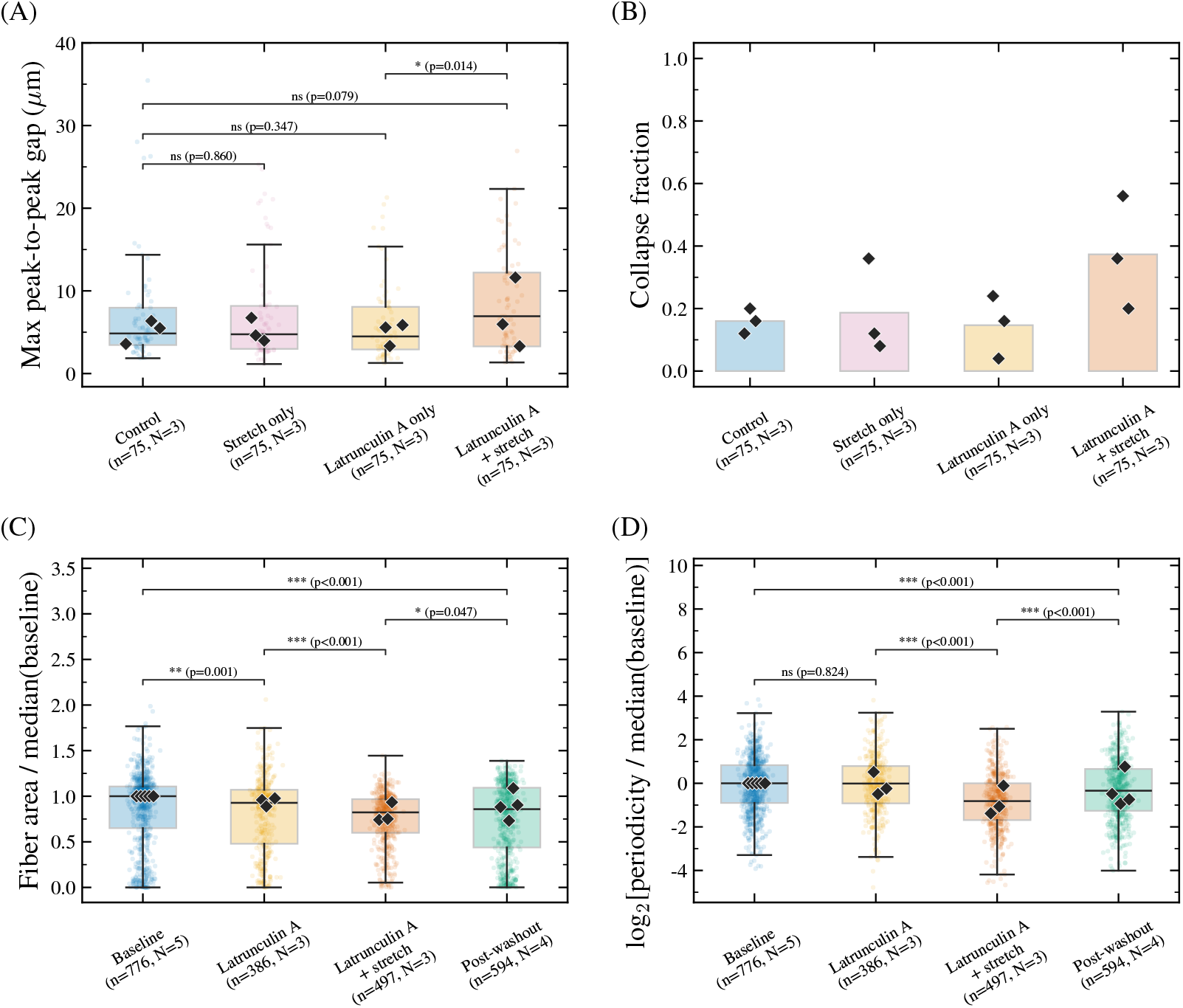
Line-ROI and endpoint analyses quantify Latrunculin A- and stretch-dependent destabilization of actin-bundle organization. **(A)** Maximum adjacent peak-to-peak gap, *G*_max_, measured from line ROIs drawn along visible GFP–*α*-actinin-positive bundle-like structures. **(B)** Line-scan collapsed ROI fraction. Detected fiber area normalized by the corresponding within-experiment baseline median. **(D)** Periodicity, shown as a log_2_ ratio relative to the corresponding within-experiment baseline median. In **(A)** and **(B)**, conditions are Control, Stretch only, Latrunculin A only, and Latrunculin A + stretch. In **(C)** and **(D)**, baseline denotes pre-Latrunculin A frames used for within-experiment normalization. Box plots show ROI-level distributions in **(A)** and tile-level distributions in **(C**,**D)**; points indicate individual ROIs or accepted tiles, and black diamonds indicate experiment-level medians. The number of line ROIs or accepted tiles *n* and independent experiments *N* are indicated for each condition. Statistical comparisons were performed using two-sided Mann–Whitney U tests for pooled ROI-level or tile-level distributions in **(A), (C)**, and **(D)** and are reported as distribution-level comparisons of technical observations.

We then used automated tile-based endpoint analysis to quantify actin-bundle organization more broadly across independent time-lapse experiments (Fig. 2C and D). Latrunculin A alone modestly reduced detected fiber area in the pooled tile-level end-point analysis, with little change in periodicity. Stretch after Latrunculin A treatment decreased both metrics, consistent with destabilization of actin-bundle organization. The pooled post-washout endpoint showed partial recovery. Together, these results show that GFP–*α*-actinin-positive organization was partly retained after Latrunculin A treatment but destabilized by stretch. We next used a stochastic filament bundle model to examine this stretch-dependent instability.

### 2.2 Modeling bundle-connectivity states

#### 2.2.1 Overview of the stochastic filament bundle model

We constructed a minimal one-dimensional stochastic model of an actin filament bundle (Fig. 3A). The model treats actin filaments as rigid intervals arranged along a one-dimensional bundle axis, with filament length set by 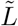. *α*-actinin is represented as an actin crosslinking protein (ACP) that forms stochastic links between actin filament pairs, with overlap-dependent binding and force-dependent catch–slip-like unbinding [26–28]. External stretch changes filament positions and modulates filament overlap. Myosin-driven contraction is implemented as contractile shortening, increasing tensile loading on bound ACPs. Representative snapshots of the collapse condition illustrate the loss of anchor-to-anchor spanning connectivity after stretch (Fig. 3B). We explored spanning connectivity across the (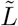, *ε*_step_) plane to examine the effects of filament length and step strain. Detailed model definitions are given in Sec. 4.6.

**Fig. 3.**
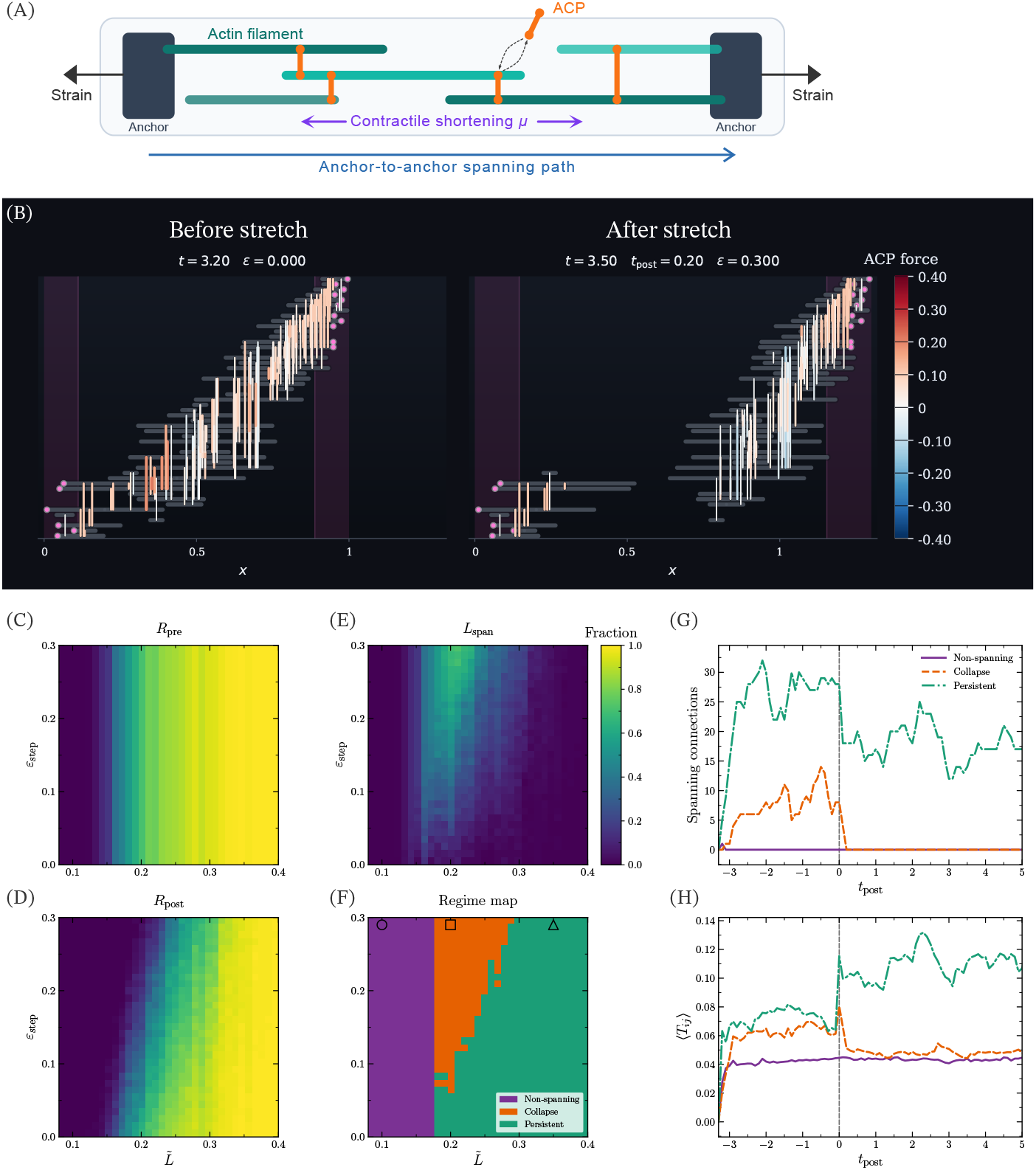
The stochastic filament bundle model generates three bundle-connectivity states across filament length and step strain. **(A)** Schematic of the stochastic filament bundle model, showing one-dimensional actin filaments, stochastic ACP links, external stretch, contractile shortening, and the anchor-to-anchor spanning path used for state classification. **(B)** Representative snapshots of the collapse condition before and after stretch, illustrating loss of anchor-to-anchor spanning connectivity. Colors indicate ACP force; red and blue denote tensile and compressive forces, respectively. **(C–E)** Ensemble-averaged heatmaps of *R*_pre_ **(C)**, *R*_post_ **(D)**, and *L*_span_ = *R*_pre_ − *R*_post_ **(E)** across 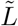 and *ε*_step_. **(F)** Ensemble-averaged regime map classifying each condition as non-spanning, collapse, or persistent; markers denote the conditions used in **(G–H)** (circle, non-spanning; square, collapse; triangle, persistent). **(G–H)** Representative trajectories corresponding to Supplementary Videos S2–S4, showing spanning connections **(G)** and mean tensile ACP load averaged over bound ACPs **(H)** for the marked conditions. The gray dashed line marks the step-strain input at *t*_post_ = 0; *t*_post_ *<* 0 denotes the pre-step response.

#### 2.2.2 Non-spanning, collapse, and persistent states

We performed ensemble simulations of the stochastic filament bundle model across the 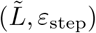 plane and quantified spanning connectivity before and after stretch (Fig. 3C–E). Here, *R*_pre_, *R*_post_, and *L*_span_ = *R*_pre_ − *R*_post_ denote pre-stretch spanning probability, post-stretch spanning probability, and stretch-induced spanning loss, respectively. Short 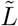 produced low *R*_pre_, whereas large 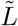 maintained high *R*_post_. Intermediate 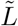 retained pre-stretch spanning but showed reduced *R*_post_ and increased *L*_span_ after stretch.

Applying the threshold classification to *R*_pre_ and *L*_span_, we assigned each condition to one of three states (Fig. 3F). Short 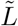 corresponded to the non-spanning state, large 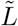 to the persistent state, and intermediate 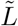 to the collapse state, in which spanning connectivity was present before stretch but lost after stretch. The collapse state occupied an extended band in the 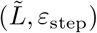 plane.

Representative trajectories and Supplementary Videos S2–S4 showed time courses of spanning connectivity and tensile ACP loading in the three states (Fig. 3G and H). Spanning connections remained near zero in the non-spanning condition, with little tensile load on bound ACPs. The collapse trajectory formed spanning connections before the strain input but lost them after stretch, accompanied by a decrease in mean tensile ACP load. The persistent trajectory maintained spanning connections and showed an increase in mean tensile ACP load after the strain input, consistent with force-supported ACP retention. These simulations show that the collapse state occurs for actin filaments of intermediate length under sufficiently large step strain. We next examine how model parameters modulate bundle formation and post-stretch collapse.

### 2.3 Parameter control of formation and collapse

#### 2.3.1 Key parameters shift the three-regime map

We varied force sensitivity, ACP binding rate, and contractile shortening to examine how these parameters shift the three-state map (Fig. 4). Across representative values of *η*_*F*_, *κ*_on_, and *µ*, the three-state structure was retained, whereas regime boundaries shifted. Changing *η*_*F*_ produced modest changes, suggesting that force sensitivity modulates regime boundaries without dominating the overall pattern. Reducing *κ*_on_ enlarged the non-spanning region and shifted persistent conditions toward collapse, indicating that ACP binding contributes to both pre-stretch formation and post-stretch maintenance. Increasing *µ* expanded the non-spanning region and shifted the collapse band, indicating that contractile shortening affects both formation and stretch-induced loss. We therefore analyzed pre-stretch formation and post-stretch loss separately.

**Fig. 4.**
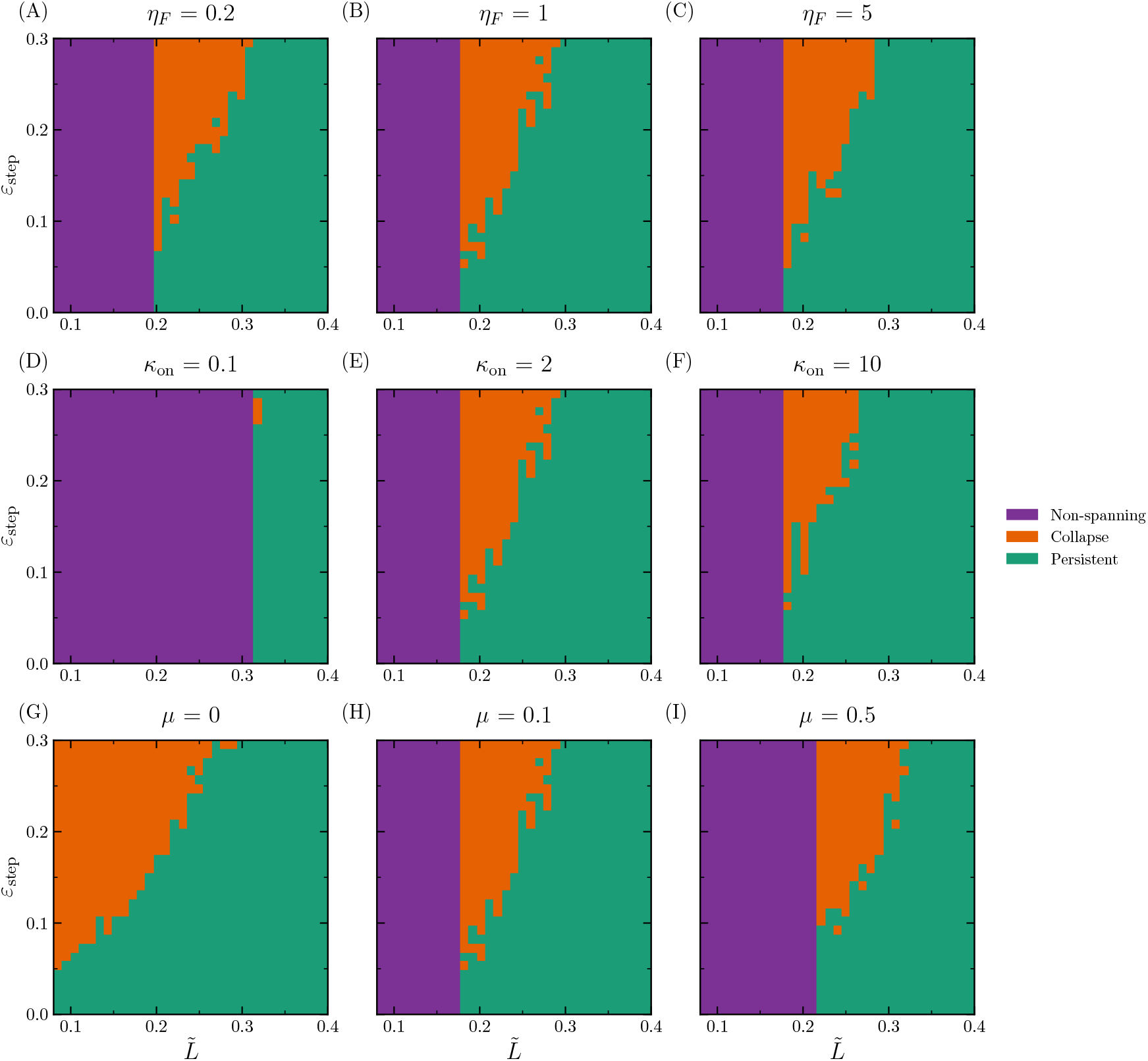
Force sensitivity, ACP binding rate, and contractile shortening shift the three-state regime map. Regime maps in the (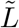, *ε*_step_) plane for representative values of force-sensitivity scale *η*_*F*_ **(A–C)**: 0.2, 1.0, 5.0; ACP binding rate *κ*_on_ **(D– F)**: 0.1, 2.0, 10.0; and actomyosin-induced contractile shortening *µ* **(G–I)**: 0, 0.1, 0.5. Colors indicate non-spanning, collapse, and persistent states assigned by the ensemble-averaged threshold rule. Full sweeps are shown in Supplementary Fig. S1.

#### 2.3.2 Controls of pre-stretch bundle formation

We grouped collapse and persistent conditions as formed states because both retain spanning connectivity before stretch. For each parameter value, *A*_formed_ and *A*_non_ denote the fractions of the 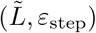 grid area assigned to formed and non-spanning states, respectively. Here, *A*_collapse_ and *A*_persistent_ denote the grid-area fractions classified as collapse and persistent, respectively:

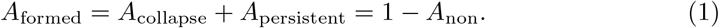

We also examined *R*_pre_ as a function of 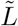 to assess local spanning formation directly (Fig. 5).

**Fig. 5.**
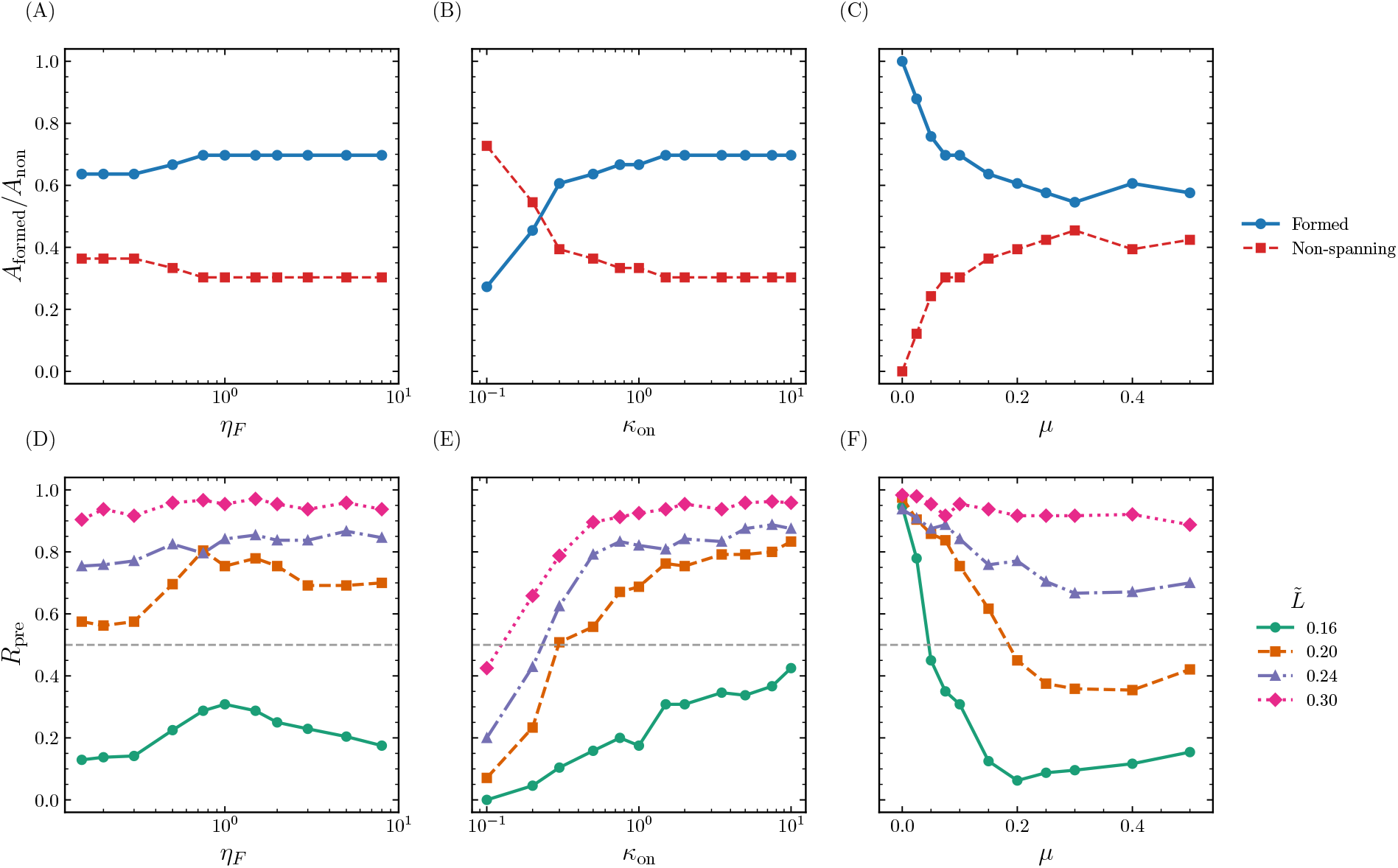
Spanning probability analysis characterizes the formed–non-spanning split before stretch. **(A–C)** Area fractions of formed states, *A*_formed_, and non-spanning states, *A*_non_, across *η*_*F*_ **(A)**, *κ*_on_ **(B)**, and *µ* **(C)**; formed states combine collapse and persistent states. **(D–F)** Pre-stretch spanning probability *R*_pre_ versus 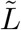 for the same parameter sweeps. The dashed line marks the formation threshold *θ*_pre_ = 0.5.

Changing *η*_*F*_ produced modest changes in both the area fractions and *R*_pre_ (Fig. 5A and D). Increasing *κ*_on_ shifted the system toward formed states and increased prestretch spanning (Fig. 5B and E). Larger *µ* expanded the non-spanning fraction and suppressed *R*_pre_ (Fig. 5C and F). Thus, pre-stretch formation depends mainly on ACP binding rate *κ*_on_ and contractile shortening *µ*, with a weaker contribution from force sensitivity *η*_*F*_ .

#### 2.3.3 Controls of post-stretch collapse

We next analyzed post-stretch loss of spanning connectivity within formed states. The fraction of formed conditions classified as collapse was defined as

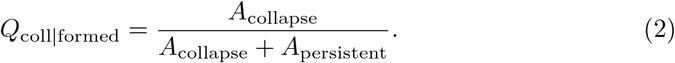

Here, *Q*_coll|formed_ = 0 means that all formed conditions are persistent, whereas *Q*_coll|formed_ = 1 indicates that all formed conditions collapse. Post-stretch loss normalized by pre-stretch spanning was quantified as

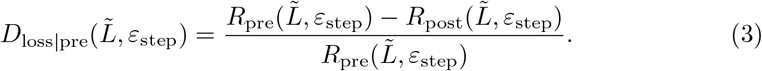

This quantity was evaluated only for formed conditions, where *D*_loss|pre_ = 0 corresponds to full retention and *D*_loss|pre_ = 1 to complete loss of pre-stretch spanning.

Changing *η*_*F*_ had a limited effect on both *Q*_coll|formed_ and *D*_loss|pre_ (Fig. 6A–D). Increasing *κ*_on_ reduced both measures, indicating that stronger ACP binding stabilizes formed bundles after stretch (Fig. 6E–H). Larger *µ* increased *D*_loss|pre_, indicating that contractile shortening enhances stretch-induced loss (Fig. 6I–L). Across formed conditions, relative loss increased with *ε*_step_ and was larger at shorter 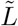. Thus, post-stretch collapse is shaped primarily by ACP binding rate and contractile shortening, with a weaker contribution from force sensitivity.

**Fig. 6.**
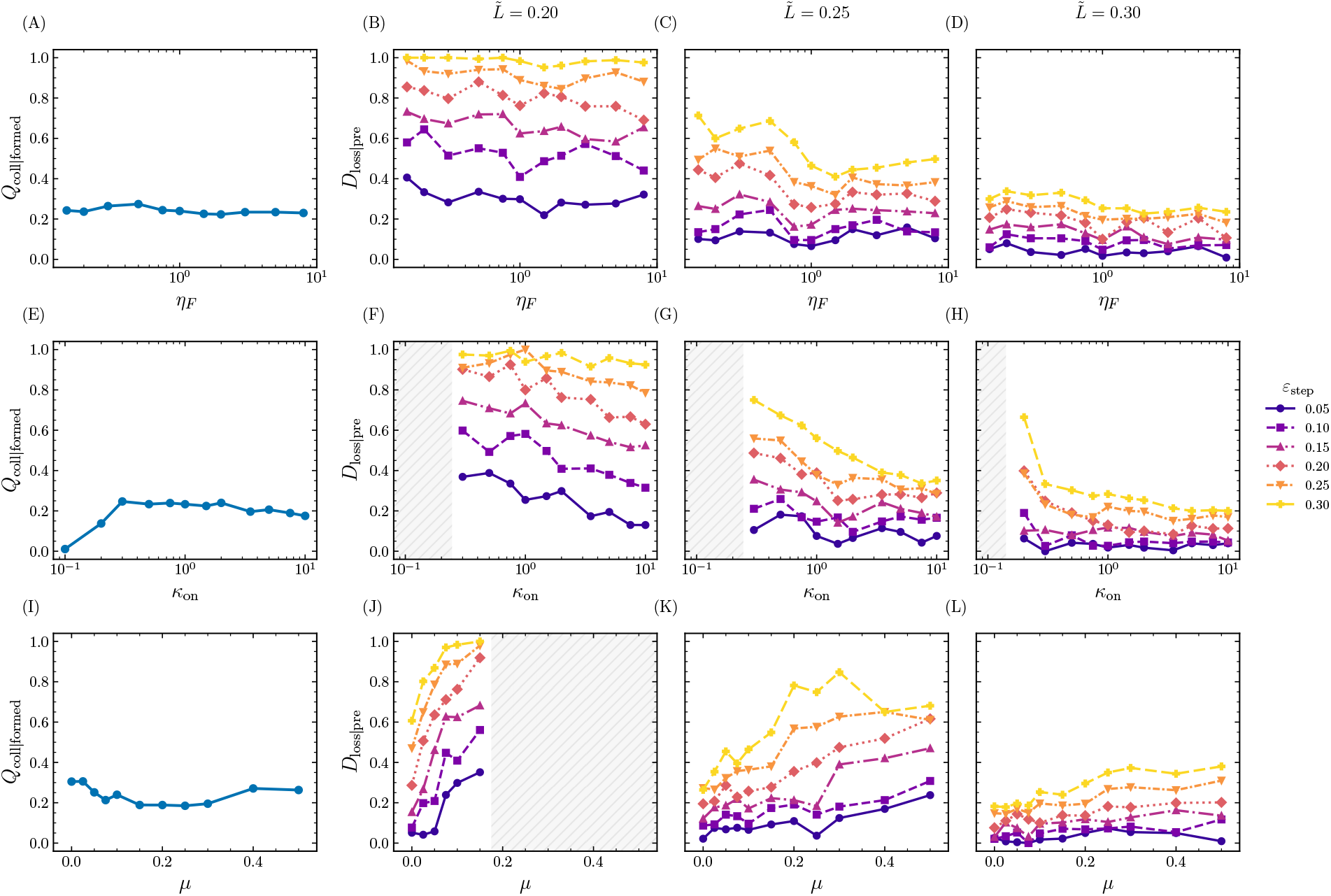
Post-stretch connectivity loss quantifies the persistent–collapse split within formed bundles. Columns show sweeps over *η*_*F*_ **(A–D)**, *κ*_on_ **(E–H)**, and *µ* **(I–L). (A**,**E**,**I)** Fraction of formed conditions classified as collapse, *Q*_coll|formed_. **(B–D**,**F–H**,**J–L)** Post-stretch loss normalized by pre-stretch spanning, *D*_loss|pre_, for selected filament lengths 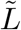. Colors, line styles, and markers indicate *ε*_step_. Gray hatched regions indicate pre-stretch non-spanning conditions excluded from the loss analysis.

These analyses separate the three-state map into pre-stretch formation and post-stretch loss. Stronger ACP binding promotes formation and post-stretch maintenance, while contractile shortening suppresses formation and increases post-stretch loss. Post-stretch collapse is most evident under larger step strain and in bundles composed of shorter actin filaments; force sensitivity has weaker effects on both formation and collapse. We next constructed a reduced model that retains four coarse controls: filament length-dependent connectivity, ACP binding rate, stretch-induced loss, and contractile shortening.

### 2.4 Reduced model of growth–loss balance

We constructed a reduced model that incorporates the four controls identified above: filament length, ACP binding, stretch-induced loss, and contractile shortening, and represents spanning connectivity by a single variable *r*. In this model, steady-state connectivity *r*^*^ follows a growth–loss balance. Filament length and ACP binding promote connectivity growth, while external stretch and contractile shortening drive connectivity loss. Evaluating *r*^*^ at *ε* = *ε*_step_ produced a connectivity landscape and, after thresholding, a three-state regime map that captured the overall stochastic organization (Fig. 7A and B). Detailed formulation and analysis are provided in Sec. 4.9.

**Fig. 7.**
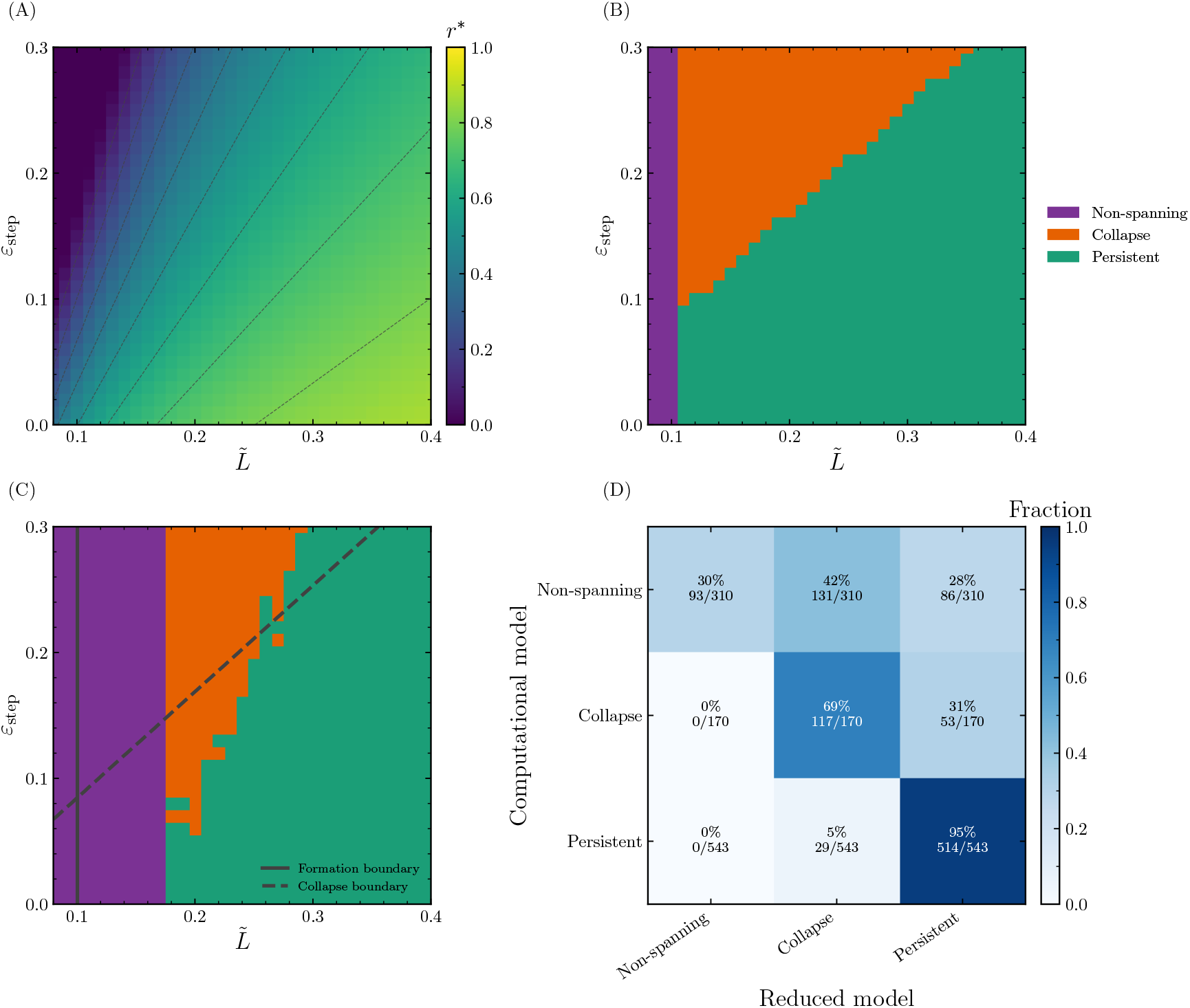
The reduced model captures the three-state regime structure of the stochastic filament bundle model. **(A)** Fixed point *r*^*^ evaluated at *ε* = *ε*_step_, shown across filament length 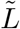 and step strain *ε*_step_. Dashed contours indicate constant *r*^*^. **(B)** Regime map obtained by applying the same threshold rule used for the stochastic model to the reduced fixed points. **(C)** Regime map of the stochastic model overlaid with formation and collapse–persistent boundaries derived from the reduced model. **(D)** State correspondence between the stochastic and reduced models, normalized within each state of the stochastic model. Colors indicate fractions, and cell labels show fractions with grid-point count ratios.

Formation and collapse–persistent boundaries from the reduced model broadly aligned with the stochastic regime map (Fig. 7C). The state correspondence matrix showed semi-quantitative agreement between stochastic and reduced state allocations (Fig. 7D). Analogous analyses in the reduced model are shown for the *κ*_on_ and *µ* sweeps in Supplementary Figs. S2–S4. This reduction summarizes the stochastic regime organization as a semi-quantitative growth–loss balance linking filament length, ACP binding, stretch-induced loss, and contractile shortening. Additional analytical consequences, including collapse-window width, critical ACP binding rate, and relaxation time, are provided in Supplementary Note 1.

## 3 Discussion

Although cellular responses to mechanical stretch have been widely studied, this study provides, to our knowledge, the first clear observations that mechanical stretching can physically rupture intracellular actin bundles. Specifically, we show that Latrunculin A-mediated inhibition of actin polymerization sensitizes intracellular actin-bundle organization to stretch in vascular smooth muscle cells. In live-cell imaging, GFP– *α*-actinin-positive organization partly persisted after Latrunculin A treatment. After mechanical stretch, detected actin-bundle coverage and periodicity both decreased, indicating destabilization of this organization. The stochastic and reduced models explained this response as a connectivity-loss mechanism in which formed bundle connectivity fails to persist after stretch.

Previous studies have linked actin polymerization to contractile and mechanical function in vascular smooth muscle cells and have used Latrunculin A to perturb actin-bundle organization and dynamics [24, 25, 29, 30]. Mechanical stretch is also known to remodel vascular smooth muscle cytoskeletal organization [31, 32]. However, the combined perturbation effects of actin polymerization inhibition and mechanical stretch have been less well defined at the level of intracellular actin-bundle organization. Our experiments show that actin-bundle organization persisting after Latrunculin A treatment is destabilized by subsequent stretch. Partial recovery after washout suggests reversible cytoskeletal destabilization, not mechanical rupture of the cell body.

A key implication of the stochastic filament bundle model is that bundle formation and mechanical retention are separable requirements. The non-spanning state represents failure to form spanning connectivity before stretch, whereas the collapse state reflects loss of already formed connectivity after stretch. This distinction explains why visible actin-bundle organization after Latrunculin A treatment does not necessarily imply mechanical robustness as a load-bearing structure. Parameter sweeps further point to ACP binding and contractile shortening as potential control points for mechanical fragility.

The reduced model identifies a growth–loss balance underlying the stochastic simulations. In this balance, filament length and ACP binding promote connectivity growth, while stretch and contractile shortening drive connectivity loss. This formulation provides a simplified framework for interpreting how changes in filament length, ACP binding, stretch, and contractile shortening shift bundle connectivity states (Supplementary Note 1). Thus, the reduced model not only explains the stochastic regime organization but also frames intracellular mechanical fragility as a growth–loss imbalance in bundle connectivity.

Disease-associated ACTA2 mutations can impair actin assembly and filament stability, suggesting that polymerization defects may affect smooth muscle mechanics at the intracellular cytoskeletal level [14–16]. Although Latrunculin A is not a genetic model of ACTA2 mutation, its use allows us to isolate the specific effect of reduced actin polymerization from the broader cellular changes associated with ACTA2 mutations. Our results suggest that reduced actin polymerization can destabilize force-bearing actin-bundle organization under tensile loading.

Future evaluation of the proposed mechanism will require measurements of actin filament-length distribution, ACP turnover, and intracellular tension in ACTA2-mutant smooth muscle cells. Integrating these measurements with spatially resolved cellular models will connect the stretch-sensitive intracellular fragility identified here to mutation-specific mechanisms of smooth muscle actinopathy [10–13]. The present study provides a cellular-scale basis for understanding how reduced actin polymerization contributes to mechanical fragility in vascular smooth muscle cells.

## 4 Materials and Methods

### 4.1 Cell culture

Rat aortic smooth muscle cells (A7r5, ATCC) were cultured with Dulbecco’s Modified Eagle Medium (1.0 g*/*L glucose; Wako) supplemented with 10% (v/v) heat-inactivated fetal bovine serum (SAFC Biosciences) and 1% penicillin–streptomycin (Wako) in a humidified 5% CO_2_ incubator at 37 ^◦^C. For experiments, cells were seeded at low density on a custom-made 20 × 20 mm^2^ stretch chamber made of polydimethylsiloxane (PDMS, 10:1 base-to-curing-agent weight ratio; SYLGARD 184, Dow Corning Toray) [33]. Before cell seeding, the bottom sheet of the chamber was coated with 10 *µ*g*/*ml bovine fibronectin (Sigma-Aldrich).

### 4.2 Plasmid and transfection

Cells were transfected with plasmids encoding GFP–*α*-actinin-1 variants (gifts from Michihiro Imamura, National Center of Neurology and Psychiatry, Japan) 24 h after seeding using Lipofectamine LTX and PLUS Reagent (Thermo Fisher Scientific) according to the manufacturer’s instructions.

### 4.3 Stretch experiment

Cells were observed using a custom-made uniaxial stretch device (STB-195, Strex) mounted on an inverted microscope (IX-71, Olympus) in a humidified 5% CO_2_ stage incubator at 37 ^◦^C. The stretch chamber consisted of a PDMS bottom membrane (∼ 90 *µ*m) supported by a cover glass (No. 000, ∼ 50 *µ*m; Matsunami Glass). Epifluorescence images were acquired with an objective lens (60 ×, NA 1.35, oil) and camera (ORCA-R2, Hamamatsu). Cells showing GFP–*α*-actinin-positive actin bundles aligned approximately parallel to the stretch axis were selected for live imaging. Because no apparent differences in actin-bundle organization were observed between the GFP–*α*-actinin-1 variants, the data were analyzed together. Before time-lapse analysis, cells were first examined under four perturbation conditions: static vehicle control, step stretch alone, Latrunculin A treatment alone, and Latrunculin A treatment followed by step stretch. Cells were observed for at least 10 min under each condition. In these observations, step stretch alone (∼ 35% strain, 2 s) did not induce apparent disruption of GFP–*α*-actinin-positive actin bundles, whereas step stretch applied in the presence of Latrunculin A (200 nM in DMSO) induced intracellular actin-bundle disruption without mechanical rupture of the cell body. Based on these observations, live-cell time-lapse imaging was performed to follow the temporal sequence of Latrunculin A treatment, stretch-induced intracellular disruption, washout-associated recovery, and mechanical release.

After baseline imaging for ∼ 40 min, Latrunculin A was added, and cells were imaged for ∼ 10 min before uniaxial step stretch was applied. Because stretching displaced the imaging field, the same cells were relocated within 1 min and imaged for an additional ∼ 10 min. Latrunculin A was washed out within 3 min by replacing the medium at least three times with fresh medium lacking Latrunculin A, followed by imaging for ∼ 40 min. Cells were then treated again with Latrunculin A for ∼ 10 min, followed by a 2 s step release to the original chamber length. After relocation within 3 min, imaging was continued for ∼ 40 min.

### 4.4 Image analysis

#### 4.4.1 Image preprocessing and cell masking

Image analysis was performed on GFP–*α*-actinin time-lapse stacks. All length parameters were applied in physical units using the spatial calibration in the TIFF metadata. For each frame, the raw intensity image *I* was robustly normalized using the frame-wise 1st and 99.5th intensity percentiles:

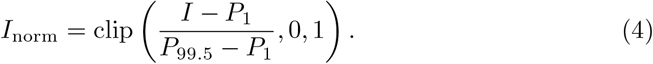

A high-pass image was obtained by subtracting a Gaussian-smoothed background with *σ* = 5 *µ*m:

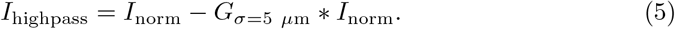

Cell masks were generated from *I*_norm_. The normalized image was smoothed with a Gaussian filter of *σ* = 2 *µ*m, and an Otsu threshold was computed from the smoothed image. A cell-core mask was defined as the largest connected component above 0.45 times the Otsu threshold. This core mask was dilated by 14 *µ*m, and the largest connected component above 0.20 times the Otsu threshold within the dilated region was used as the cell foreground. The final foreground mask was eroded by 0.3 *µ*m. During connected-component extraction, masks were hole-filled and morphologically cleaned by binary closing and opening.

#### 4.4.2 Detection of GFP–*α*-actinin-positive bundle mask

The bundle mask was detected without manual regions of interest. A multi-scale white top-hat response was computed using square structuring elements with characteristic half-widths corresponding to 0.22, 0.35, 0.55, and 0.80 *µ*m. The maximum response across scales was then robustly scaled to the range 0 to 1 using the 1^st^ and 99^th^ percentiles within the cell foreground. Pixels were assigned to the detected bundle mask when they satisfied all of the following conditions: they were inside the foreground mask, the top-hat score was above the foreground 48^th^ percentile, the top-hat score exceeded the local mean plus 0.01 times the local standard deviation, *I*_norm_ was above the foreground 8^th^ percentile, and *I*_highpass_ was above the foreground 2^nd^ percentile. Local mean and standard deviation were computed using a 1.6 *µ*m square window. After thresholding, binary opening was applied once, and small low-signal connected components were removed. Finally, connected components with area below 0.02 *µ*m^2^ were excluded from the final bundle mask. The same preprocessing, masking, and detection parameters were applied to all frames and all experimental conditions, and mask and detection quality were verified using overlay images.

#### 4.4.3 Tile-based quantification of fiber area and periodicity

The cell foreground was analyzed using square tiles of 10 *µ*m × 10 *µ*m with a stride of 5 *µ*m. Tiles with foreground fraction below 0.45 were excluded. Each accepted tile was treated as one technical observation.

Detected fiber area was defined as the fraction of pixels in an accepted tile assigned to the GFP–*α*-actinin-positive bundle mask:

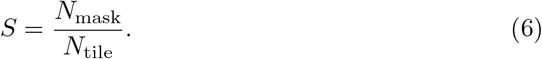

Here, *N*_tile_ denotes the total number of pixels in the accepted tile. For figure display, *S* was normalized by the median value of the corresponding baseline tiles from the same experiment:

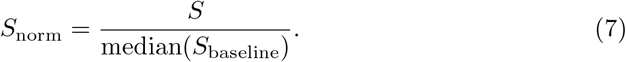

Periodicity was computed from the high-pass image in each accepted tile. Pixels outside the foreground mask were filled with the mean intensity of foreground pixels within the tile. The tile mean was then subtracted, a two-dimensional Hann window was applied, and the Fourier power spectrum was computed:

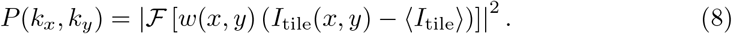

Here, *I*_tile_ denotes the high-pass tile image after foreground filling. The radial spatial frequency was defined as

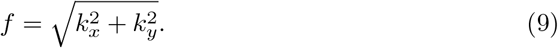

The periodicity score was defined as the ratio of power in the *α*-actinin periodic band, 0.4 ≤ *f* ≤ 1.0 cycles*/µ*m, to power in the low-frequency band, 0.15 ≤ *f <* 0.4 cycles*/µ*m:

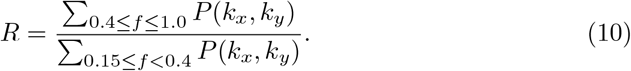

For figure display, periodicity was normalized by the median value of the corresponding baseline tiles from the same experiment and plotted on a log_2_ scale:

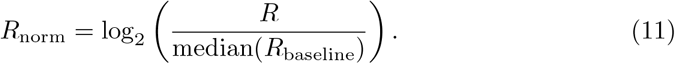

For the representative time course, each metric was normalized to the median of all accepted pre-Latrunculin A control tiles from the same experiment; time-course traces show the tile median with the interquartile range.

#### 4.4.4 Line-ROI validation of local bundle collapse

To quantify local discontinuity of GFP–*α*-actinin-positive bundle organization, we performed manual line-ROI analysis under four conditions: untreated control (Control), stretch without Latrunculin A (Stretch only), Latrunculin A without mechanical stretch at imaging (Latrunculin A only), and stretch after Latrunculin A treatment (Latrunculin A + stretch). For each condition, 25 line ROIs were manually drawn in each of three representative frames along visible GFP–*α*-actinin-positive bundle-like structures, giving *n* = 75 ROIs per condition. ROIs had a width of 5 pixels, corresponding to approximately 0.32 *µ*m, and variable lengths to follow local bundle-like structures; intensity profiles were extracted from the corresponding raw TIFF images. Frames were selected at approximately matched post-perturbation times: 10– 14 min after stretch for Stretch only, 10–12 min after Latrunculin A addition for Latrunculin A only, and 10–13.5 min after Latrunculin A addition for Latrunculin A + stretch. For the combined perturbation, frame selection was based primarily on time after Latrunculin A addition because the timing and duration of stretch varied among movies. For each line ROI, the raw intensity profile was denoted by *I*(*s*), where *s* is the distance along the ROI. The following preprocessing was applied only to one-dimensional line profiles for peak detection and was independent of the two-dimensional high-pass image used for tile-based analysis. To detect local *α*-actinin peaks after removing slow background variation, a high-pass profile was obtained by subtracting a Gaussian-smoothed background with *σ* = 3.0 *µ*m:

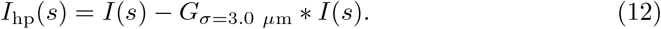

To suppress high-frequency noise, the high-pass profile was then smoothed with a narrower Gaussian filter:

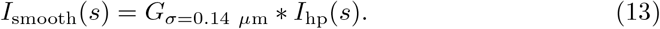

The smoothed profile was then converted to a robust *z*-score:

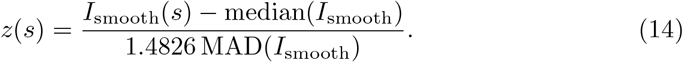

Here, MAD denotes the median absolute deviation, and the factor 1.4826 rescales MAD to estimate the standard deviation for a normally distributed signal. This normalization was used only for peak detection; the reported gap metric was measured in micrometers from the detected peak positions.

Peaks were detected on *z*(*s*) using `scipy.signal.find peaks`. The minimum peak distance was set to 0.55 *µ*m, and the prominence threshold was

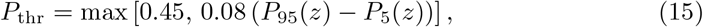

where *P*_95_(*z*) and *P*_5_(*z*) denote the 95th and 5th percentiles of each ROI-level *z*(*s*) profile, respectively. To avoid detecting spurious peaks in weak-signal regions, candidate peaks were retained only when the smoothed high-pass intensity and raw intensity at the peak position exceeded the 50th and 40th percentiles, respectively, of all line-profile points in the same frame.

For each ROI, adjacent peak-to-peak gaps *d*_*i*_ were computed from the detected peak positions *p*_*i*_, and their maximum was defined as *G*_max_:

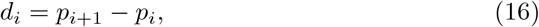

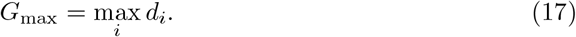

If fewer than two peaks were detected, *G*_max_ was set to the ROI length *L*_ROI_. ROIs with *G*_max_ ≥ 10 *µ*m were classified as collapsed, and their fraction was reported as the line-scan collapsed ROI fraction. This collapse threshold was set conservatively above the typical adjacent peak-to-peak gaps in control line scans, which had a median of 1.73 *µ*m and a 95th percentile of 5.25 *µ*m.

### 4.5 Statistical analysis

For the line-ROI validation analysis in Fig. 2A, ROI-level distributions of the maximum peak-to-peak gap, *G*_max_, were compared using two-sided Mann–Whitney U tests. These tests used pooled ROI-level observations and were treated as distribution-level comparisons, with experiment-level medians shown separately. The line-scan collapsed ROI fraction in Fig. 2B was computed from the same *G*_max_ values and shown descriptively without statistical testing.

Endpoint analysis for Fig. 2C and D used the first stimulation cycle from each independent experiment, with one representative frame selected for each condition within each experiment. Conditions were grouped as Baseline, Latrunculin A, Latrunculin A + stretch, and post-washout. The Baseline group included pre-Latrunculin A frames, including stretch-only pre-drug frames when present, as the pre-drug reference for actin-bundle organization. The post-washout group included washout-only, washout + stretch, and washout + mechanical release conditions. For each condition present in an experiment, the endpoint frame was defined as the final frame of the first continuous interval assigned to that condition. Frames annotated as quality caution, dying, saturated, or overexposed were excluded. For baseline-normalized analyses, each experiment was normalized to its own baseline-tile median; experiments without a baseline endpoint were excluded from these analyses. For the tile-based endpoint analysis, *n* denotes the number of accepted tiles and *N* denotes the number of independent time-lapse experiments. Pooled tile-level distributions were compared using two-sided Mann–Whitney U tests; these tests were treated as distribution-level comparisons of image-derived technical observations, with independent-experiment numbers reported separately.

For all Mann–Whitney U tests in Fig. 2, reported *p* values were not adjusted for multiple comparisons. Significance labels were assigned as *p* ≥ 0.05 for ns, *p <* 0.05 for *, *p <* 0.01 for **, and *p <* 0.001 for * * *.

### 4.6 Stochastic filament bundle model

#### 4.6.1 Filament geometry and force balance

We model the actin bundle as a one-dimensional array along the bundle axis. Each actin filament *i* is represented as a rigid interval of length *L*_*i*_, with initial center position 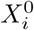. We write the current center position as

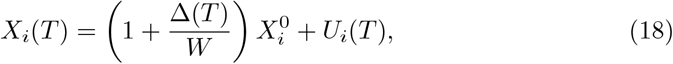

where *W* is the bundle length, Δ(*T*) is the displacement imposed by external stretch, and *U*_*i*_(*T*) is the nonaffine displacement from the imposed affine position.

Lengths are nondimensionalized by *W*, energies by *K*_*X*_*W* ^2^, and time by a characteristic ACP turnover time *T*_0_, where *K*_*X*_ is the effective ACP stiffness. Let *A* denote the dimensional anchor width described below. We define

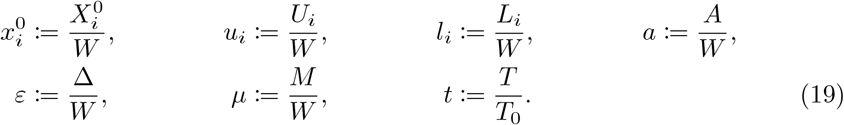

Here, *M* denotes the myosin-induced contractile shortening. All equations below use these nondimensional variables.

The nondimensional center position is

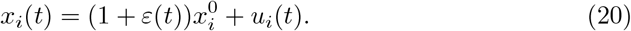

The filament endpoints are

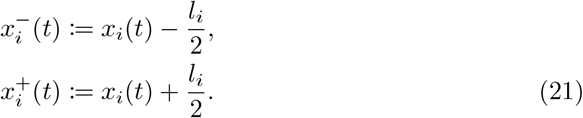

The pairwise overlap length and overlap fraction are

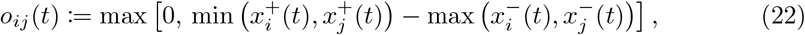

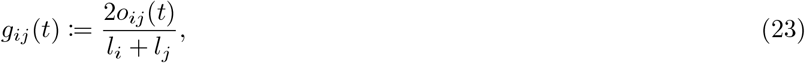

where *g*_*ij*_(*t*) measures the fractional overlap between the two actin filaments.

We define an oriented pair set 𝒫 by ordering each filament pair from left to right in the initial configuration. For each (*i, j*) ∈ 𝒫, 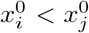. The initial center-to-center distance and signed extension are

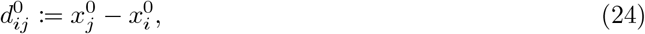

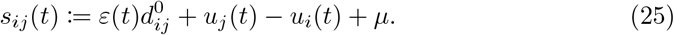

Positive *s*_*ij*_ indicates tensile extension. The parameter *µ* represents actomyosin-induced shortening of the preferred pair separation; in this formulation, positive *µ* increases the signed ACP extension generated by contraction.

We assign each pair a binary ACP state *n*_*ij*_(*t*) ∈ {0, 1}, where *n*_*ij*_ = 1 denotes an effective ACP crosslink and *n*_*ij*_ = 0 denotes no crosslink. The nondimensional elastic energy reads

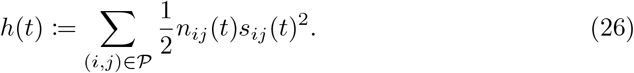

In the initial configuration, the left and right anchor sets, *B*_*L*_ and *B*_*R*_, are defined by overlap with the anchor regions [0, *a*] and [1 − *a*, 1], respectively. The anchor assignment is held fixed. Anchor filaments follow the imposed affine stretch and satisfy

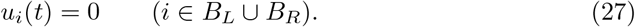

For non-anchor filaments, the overdamped dynamics obey

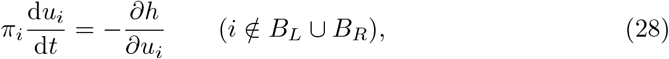

where *π*_*i*_ is the ratio of the mechanical relaxation time to the ACP turnover time. Motivated by fast mechanical relaxation in dynamically crosslinked actin networks [34– 37], we use the quasi-static limit *π*_*i*_ ≪ 1:

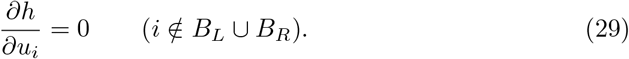

This limit is enforced by solving mechanical equilibrium after each ACP-state update.

#### 4.6.2 Stochastic crosslink dynamics

The binary ACP state *n*_*ij*_(*t*) evolves through stochastic binding and unbinding. For unbound pairs, the nondimensional binding rate is

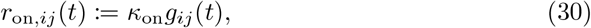

where *κ*_on_ sets the nondimensional ACP binding rate.

We define the tensile ACP load as the positive part of the signed extension,

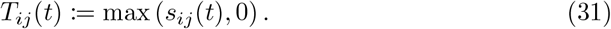

For bound pairs, the nondimensional unbinding rate follows a catch–slip form:

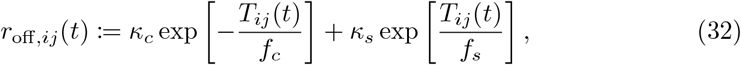

where *κ*_*c*_ and *κ*_*s*_ set the catch and slip prefactors, and *f*_*c*_ and *f*_*s*_ set the corresponding force scales. Only tensile load enters this rate; compression affects force balance but not unbinding directly. Force sensitivity is varied by scaling the catch and slip force scales with a common factor *η*_*F*_ :

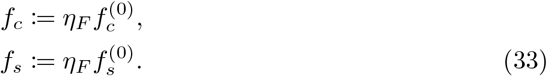

Force sensitivity increases for smaller *η*_*F*_ and decreases for larger *η*_*F*_, while the zero-load unbinding rate remains

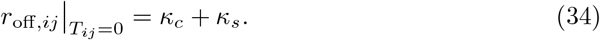

Overlap is required for an occupied ACP state:

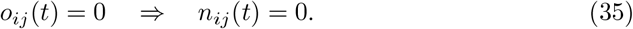

For *o*_*ij*_(*t*) *>* 0, *n*_*ij*_(*t*) follows a two-state stochastic process, with 0 and 1 denoting unbound and bound states:

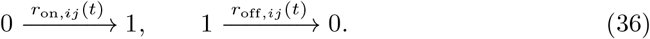

For finite time steps, these continuous-time transition rates were converted into Bernoulli transition probabilities. At each time step, an unbound pair was assigned a binding event with probability

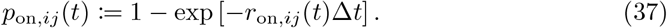

A bound pair was assigned an unbinding event with probability

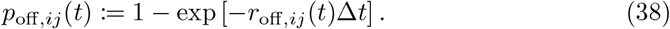

#### 4.6.3 Spanning analysis and regime classification

At each time point, we define a graph whose nodes are actin filaments and whose edges are occupied ACP links with *n*_*ij*_(*t*) = 1 and *o*_*ij*_(*t*) *>* 0. Spanning is defined as the existence of a path in this graph from the left anchor set *B*_*L*_ to the right anchor set *B*_*R*_. Spanning connections are quantified by the minimum cut between *B*_*L*_ and *B*_*R*_ in the occupied ACP graph. Each occupied ACP edge has unit capacity, so the cut value equals the minimum number of occupied ACP links required to disconnect the two anchor sets.

We compute the mean tensile ACP load as

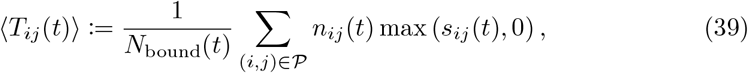

with

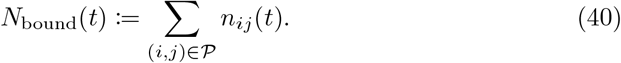

The average is taken over all bound ACPs. If *N*_bound_(*t*) = 0, we set ⟨*T*_*ij*_(*t*)⟩ = 0.

Let 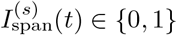 denote the spanning indicator for seed *s*. Let 𝒲_pre_ and 𝒲_post_ denote the pre- and post-stretch time windows used for averaging. For each seed, we compute pre- and post-stretch spanning probabilities as

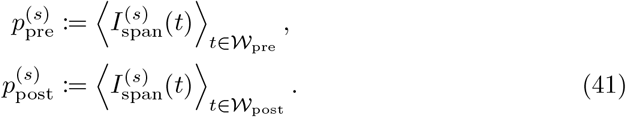

The ensemble-averaged probabilities are

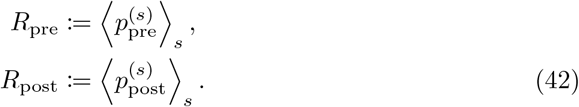

We define the stretch-induced spanning loss as

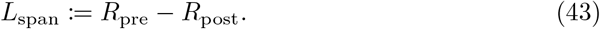

Each condition is classified using two thresholds,

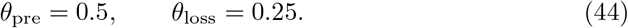

The regime label is assigned as

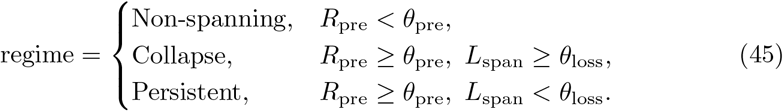

These labels distinguish pre-stretch formation failure, stretch-induced loss, and post-stretch maintenance of spanning connectivity.

### 4.7 Simulation protocol

#### 4.7.1 Initial actin filament ensemble

For each simulation, initial filament centers 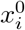 are sampled uniformly along the bundle axis subject to containment within the domain. Filament lengths are sampled from a truncated exponential distribution,

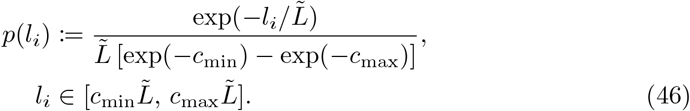

Exponential filament-length distributions are consistent with electron microscopic measurements of in vitro polymerized actin filaments and actin filament length measurements in nonmuscle actomyosin bundles [38, 39]. Here, 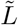 is the characteristic filament length, and *c*_min_ and *c*_max_ set the lower and upper truncation factors. The parameter *ρ*_*A*_ sets the target total filament length density, and the filament number is chosen as

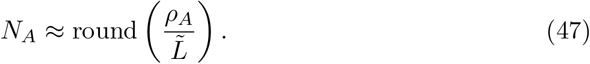

After filament generation, the filament set and the candidate ACP pair set 𝒫, comprising all filament pairs, are fixed; only the ACP states *n*_*ij*_(*t*) evolve stochastically. Binding has a nonzero probability only for pairs with positive instantaneous overlap, and bound ACPs are removed when their overlap vanishes.

#### 4.7.2 Mechanical equilibration and stochastic update

Simulations use a fixed time step Δ*t*. At each step, we first solve the quasi-static force-balance equation (Eq. (29)) for the current strain *ε*(*t*) and ACP state *n*_*ij*_(*t*). We then compute the instantaneous overlaps, overlap fractions, signed extensions, and tensile ACP loads. Bound ACPs with zero overlap are removed according to Eq. (35), and the remaining ACP states are updated by Bernoulli sampling using Eq. (37) and Eq. (38). Mechanical equilibrium is then recomputed after each ACP-state update. All candidate ACP pairs are initialized in the unbound state, *n*_*ij*_(0) = 0. The initial zero-strain phase allows stochastic ACP binding to form and equilibrate bundle connectivity before the step strain.

#### 4.7.3 Strain protocol and readouts

Simulations start with a zero-strain phase, *ε* = 0. At *t* = *t*_step_, we impose an instantaneous step strain by setting *ε* : 0 → *ε*_step_. The ACP state is not updated at the step frame; mechanical equilibrium is solved under the new strain with the same ACP configuration. The system is then evolved at fixed *ε* = *ε*_step_. Ensemble maps used Δ*t* = 0.2, whereas representative single-seed trajectories used Δ*t* = 0.1 for smoother visualization. Both protocols used the same pre-step duration, post-step duration, and total simulation time.

Pre- and post-stretch spanning probabilities are computed as time averages of the spanning indicator over finite windows. The pre-stretch window contains the final frames before the step strain, and the post-stretch window contains the final frames of the simulation. The same post-window definition is used for the no-stretch control, *ε*_step_ = 0.

### 4.8 Computational parameters and parameter sweeps

#### 4.8.1 Baseline computational parameters

Fixed parameters for the baseline simulations are listed in Table 1. We use this parameter set as a representative condition after confirming that the qualitative three-state regime structure was robust to parameter variation. The parameter sweeps below examine how key parameters shift the regime boundaries.

**Table 1.** Baseline parameters for stochastic filament bundle simulations.

| Parameter | Value | Meaning |
| --- | --- | --- |
| $\rho_A$ | 10.0 | target total filament length density |
| $a$ | 0.1 | anchor width |
| $\mu$ | 0.1 | actomyosin-induced contractile shortening |
| $\kappa_{\text{on}}$ | 2.0 | ACP binding rate coefficient |
| $\kappa_c$ | 1.5 | catch unbinding rate coefficient |
| $\kappa_s$ | 0.01 | slip unbinding rate coefficient |
| $f_c^{(0)}$ | 0.3 | baseline catch force scale |
| $f_s^{(0)}$ | 0.1 | baseline slip force scale |
| $\eta_F$ | 1.0 | force-sensitivity scaling factor |
| $c_{\text{min}}$ | 0.2 | lower filament-length truncation factor |
| $c_{\text{max}}$ | 2.5 | upper filament-length truncation factor |
| $\Delta t$ | 0.2 | time step for ensemble simulations |
| $t_{\text{step}}$ | 3.2 | step-strain time |
| $t_{\text{post}}$ | 14.4 | post-step duration |
| $t_{\text{total}}$ | 17.6 | total simulation time |
| Window size | 5 frames | pre/post spanning window size |

For the baseline ensemble map, we swept filament length and step strain over

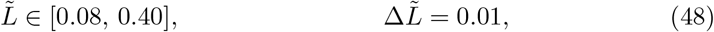

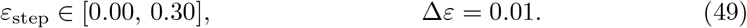

Each grid point used *N*_seeds_ = 96 independent simulations.

#### 4.8.2 Extended parameter sweeps

Extended sweeps used the same 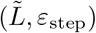 grid as the baseline ensemble map, with *N*_seeds_ = 48 independent simulations per grid point. All parameters followed Table 1 except for the swept parameter. We swept force sensitivity over *η*_*F*_ ∈ {0.15, 0.2, 0.3, 0.5, 0.75, 1.0, 1.5, 2.0, 3.0, 5.0, 8.0}, ACP binding rate over *κ*_on_ ∈ {0.1, 0.2, 0.3, 0.5, 0.75, 1.0, 1.5, 2.0, 3.5, 5.0, 7.5, 10.0}, and contractile shortening over *µ* ∈ {0, 0.025, 0.05, 0.075, 0.1, 0.15, 0.2, 0.25, 0.3, 0.4, 0.5}.

### 4.9 Reduced model

#### 4.9.1 Reduced order parameter and dynamics

The reduced model coarse-grains stochastic filament geometry, ACP turnover, force-sensitive unbinding, and actomyosin-induced contractile shortening into a scalar connectivity variable *r*(*t*) ∈ [0, 1]. Here, *r* = 0 represents a non-spanning state, *r >* 0 a spanning state, and *r* ≃ 1 a redundant spanning state.

We define the reduced dynamics as a growth–loss equation,

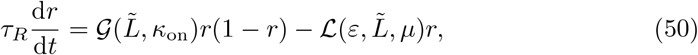

where 𝒢 is the connectivity growth rate, ℒ is the connectivity loss rate, and *τ*_*R*_ sets the timescale of changes in *r*. The growth rate depends on 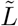 and *κ*_on_ through overlap-limited ACP rebinding. The loss rate depends on *ε*, 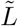, and *µ* through stretch-induced overlap loss, load sharing, and contractile loading. The growth term combines the need for an existing partial spanning connection, the remaining capacity for additional connectivity, and a rebinding-dependent growth coefficient, giving 𝒢*r*(1 − *r*). The loss term is linear in *r* and represents removal of existing spanning connectivity by stretch-and contraction-dependent loading.

The connectivity growth rate combines geometric opportunity and ACP rebinding availability:

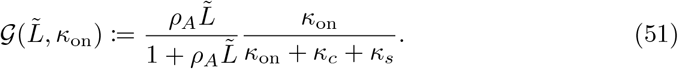

The geometric factor uses 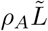 as a proxy for available overlapping actin material, with 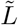 denoting the mean filament length of the exponential distribution. It saturates when geometric opportunity is no longer limiting. The rebinding factor follows the zero-load two-state ACP balance, with *κ*_on_ competing against the basal unbinding rate *κ*_*c*_ + *κ*_*s*_.

We define the connectivity loss rate to coarse-grain force-dependent ACP loss, stretch-induced overlap loss, and load sharing:

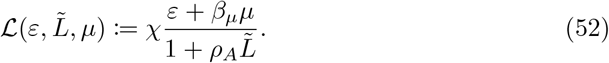

The numerator gives the effective load supported by spanning connectivity, combining external stretch with actomyosin-induced contractile shortening converted by *β*_*µ*_ into strain-equivalent load. The denominator uses 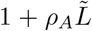 as a coarse proxy for the geometric capacity to distribute load across the filament bundle. The coefficient *χ* converts this stretch- and contraction-dependent load into connectivity loss.

#### 4.9.2 Fixed points and regime classification

For fixed 𝒢 and ℒ, the right-hand side of Eq. (50) can be written as

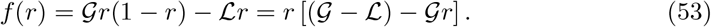

The fixed points are

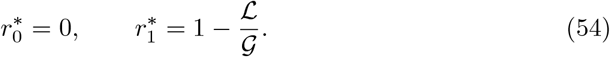

The zero fixed point 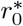 denotes the non-spanning fixed point, stable for 𝒢 ≤ ℒ, whereas the nonzero fixed point 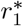 denotes the formed fixed point, stable for 𝒢 *>* ℒ. Thus, the steady-state value of the spanning connectivity is

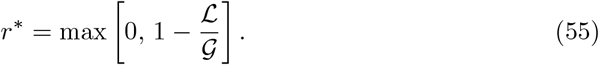

We define the growth capacity *K* and contractile-loss coefficient *δ*_*µ*_ by

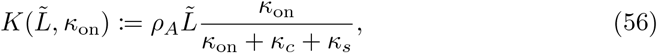

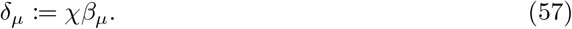

Using Eq. (51) and Eq. (52), the fixed-point connectivity can be rewritten as

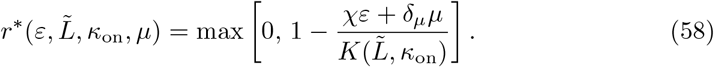

Evaluating this expression at *ε* = 0 and *ε* = *ε*_step_ gives the pre- and post-stretch values, respectively:

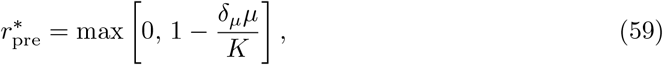

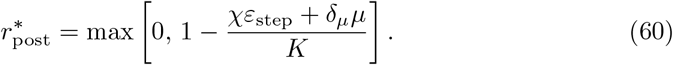

To compare with the full stochastic model, we apply the same operational thresholds as in Eq. (44). The regime label in the reduced model is assigned as

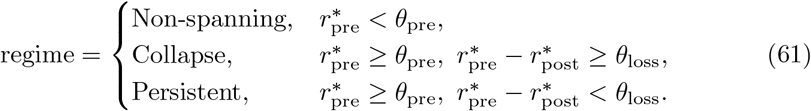

Additional analytical relations derived from the reduced model, including regime-boundary expressions, collapse-window width, critical ACP binding rate, and relaxation timescale, are provided in Supplementary Note 1.

#### 4.9.3 Calibration of parameters in the reduced model

We calibrate the parameters *δ*_*µ*_ and *χ* in the reduced model against the stochastic filament bundle model. The conversion coefficient *β*_*µ*_ is then computed as

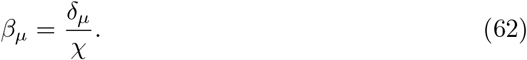

We do not calibrate *τ*_*R*_, since static regime maps do not constrain the timescale of the reduced model. Calibration uses simulation results from the stochastic filament bundle model for the *κ*_on_ and *µ* sweeps. The *η*_*F*_ sweep is excluded because *η*_*F*_ does not enter the reduced model.

For each candidate pair (*δ*_*µ*_, *χ*), we compute 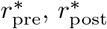 and the regime label from the reduced model using Eq. (61). We compare this label with the regime label from the stochastic filament bundle model and maximize a macro F1 score for joint three-state classification. The calibrated values are

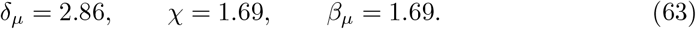

These values are fixed for all comparisons with the reduced model and are not reestimated for individual figures.

## Acknowledgements

The authors thank T.S. Matsui for advice on the experiments and for discussions on the underlying mechanisms.

## Statements and Declarations

### Ethics approval

Not applicable. This study used an established rat vascular smooth muscle cell line and did not involve human participants, human data, human biological material, or live animals.

### Consent to participate

Not applicable.

### Consent for publication

Not applicable.

### Data and code availability

Data and code are available from the corresponding authors upon reasonable request.

### Author contributions

**Eiji Matsumoto**: Conceptualization, Methodology, Formal analysis, Investigation, Data curation, Software, Visualization, Writing - original draft, Writing - review and editing, Funding acquisition. **Shinji Deguchi**: Conceptualization, Methodology, Formal analysis, Investigation, Data curation, Visualization, Supervision, Project administration, Resources, Writing - original draft, Writing - review and editing, Funding acquisition.

### Use of generative AI and AI-assisted technologies

During manuscript preparation, the authors used ChatGPT (OpenAI) to assist with English-language editing and text rephrasing. The authors reviewed and edited all AI-assisted text and take full responsibility for the final content.

## Supplementary Note 1: Analytical consequences of the reduced model

### Regime boundaries and characteristic timescales

Linearizing the reduced dynamics near the stable formed fixed point gives an analytical relaxation timescale. The corresponding linear relaxation rate is 𝒢 − ℒ, so

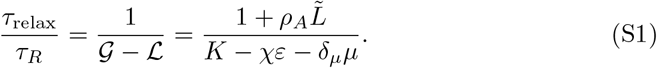

This timescale diverges as *τ*_relax_*/τ*_*R*_ → ∞ when 𝒢 − ℒ → 0^+^.

The operational thresholds define two boundaries along the 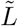 axis: a formation boundary and a collapse–persistent boundary. The formation boundary, 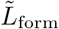, follows from 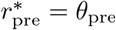:

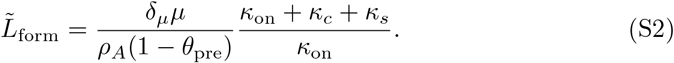

The collapse–persistent boundary, 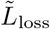, follows from 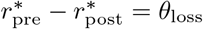:

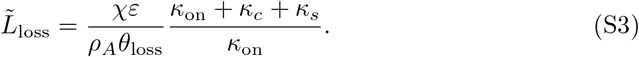

The collapse-window width along the 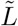 axis is

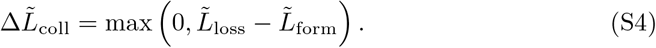

Using Eq. (S2) and Eq. (S3), this becomes

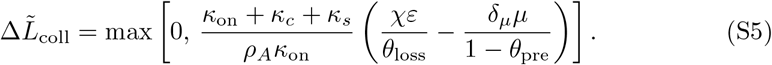

A nonzero collapse window appears when

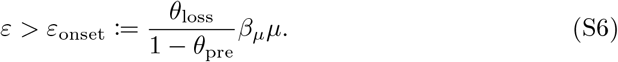

For *θ*_pre_ = 0.5 and *θ*_loss_ = 0.25, this gives

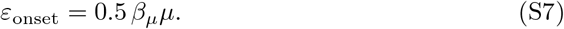

The persistent state requires the growth capacity to exceed both the formation and loss demands:

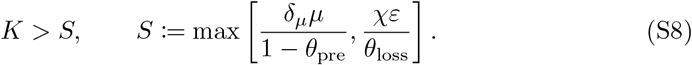

Solving this condition for *κ*_on_ gives the persistent-state boundary in ACP binding rate,

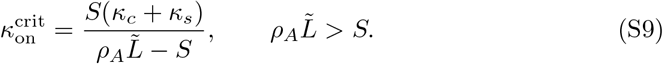

### Analytical readouts from the reduced model

Here, we use the reduced equations to examine collapse-window width, rebinding rescue, and relaxation timescales (Fig. S5).

The collapse-window width 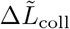 (Eq. (S4)) quantifies the range of 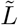 over which collapse is expected. It increased with step-strain amplitude, indicating that stronger stretch broadens the collapse window along the 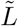 axis (Fig. S5A). The onset condition in Eq. (S6) gives the step-strain amplitude required for a nonzero collapse window.

Contractile shortening shifted this onset by changing the pre-stretch formation boundary. Together, these expressions predict both where collapse appears in parameter space and the width of the collapse window along the 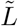 axis.

The persistent-state condition in Eq. (S8) can be solved for the minimum ACP binding rate required to maintain persistence, 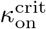 (Eq. (S9)). This critical rate increased for shorter 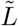 and larger *ε*_step_, showing that stronger ACP binding is needed to compensate for stretch-induced loss under weaker geometric connectivity or stronger stretch (Fig. S5B). This relation predicts the ACP binding rate required to maintain persistence under applied stretch.

Linearizing around the stable formed fixed point gives the relaxation timescale (Eq. (S1)). The resulting timescale increased near the collapse–persistent boundary, indicating slowing as the formed fixed point approached loss of stability (Fig. S5C). Representative trajectories showed the corresponding dynamics: rapid persistence far from the boundary, slow relaxation near the boundary, and collapse after loss of the formed fixed point (Fig. S5D). Thus, this timescale relation provides a coarse-grained readout of how rapidly bundle connectivity relaxes near the collapse–persistent boundary.

Together, these analytical consequences summarize the computational results as relations linking filament length, ACP binding, stretch-induced loss, and contractile shortening. These relations connect the three-state regime map to collapse-window width, rebinding rescue, and the characteristic response timescale.

## Supplementary Figures

**Fig. S1.**
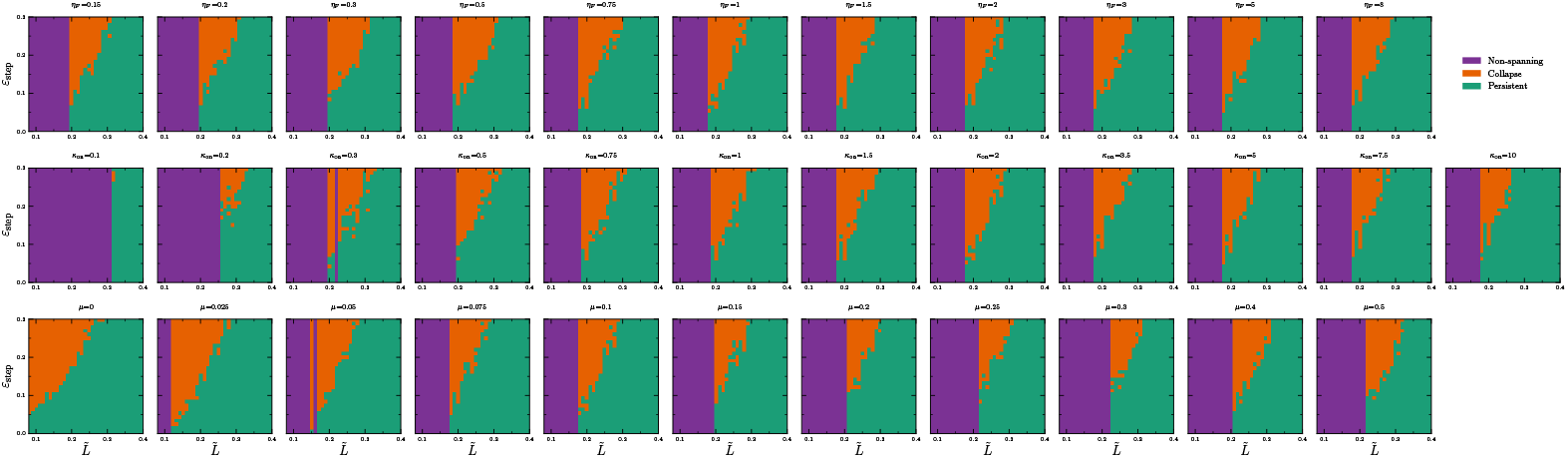
Full parameter sweeps of the stochastic filament bundle model regime map. Regime maps in the (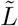, *ε*_step_) plane for all simulated values of the force-sensitivity scale *η*_*F*_ (top row), ACP binding rate *κ*_on_ (middle row), and actomyosin-induced contractile shortening *µ* (bottom row). Colors indicate non-spanning, collapse, and persistent states assigned by the ensemble-averaged threshold rule.

**Fig. S2.**
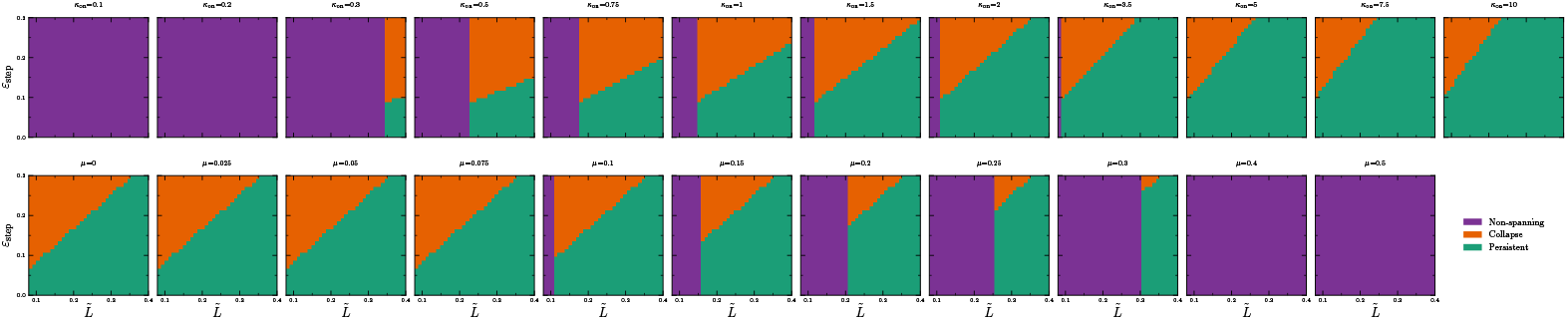
Regime maps of the reduced model across ACP binding and contractile shortening sweeps. Regime maps in the (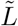, *ε*_step_) plane generated from the reduced fixed-point classification. The top row shows the ACP binding-rate sweep over *κ*_on_. The bottom row shows the actomyosin-induced contractile shortening sweep over *µ*. Colors indicate non-spanning, collapse, and persistent states assigned by the same operational threshold rule as the stochastic model.

**Fig. S3.**
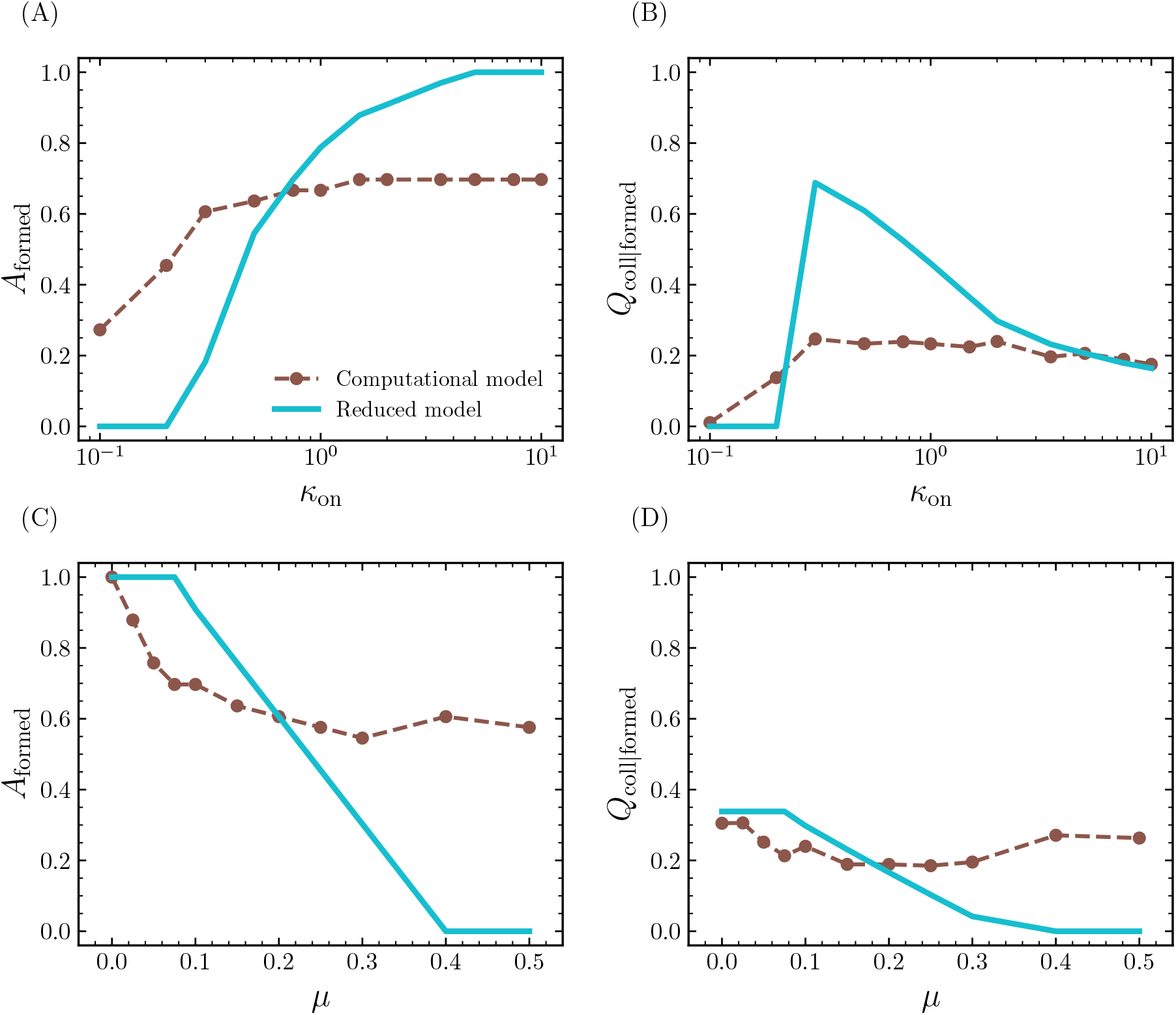
Summaries from the reduced model capture the main formation and collapse trends. Comparison between computational model and reduced model summary metrics for the ACP binding rate sweep and contractile shortening sweep. **(A**,**C)** Formed area fraction *A*_formed_ across *κ*_on_ **(A)** and *µ* **(C). (B**,**D)** Fraction of formed conditions classified as collapse, *Q*_coll|formed_, across *κ*_on_ **(B)** and *µ* **(D)**.

**Fig. S4.**
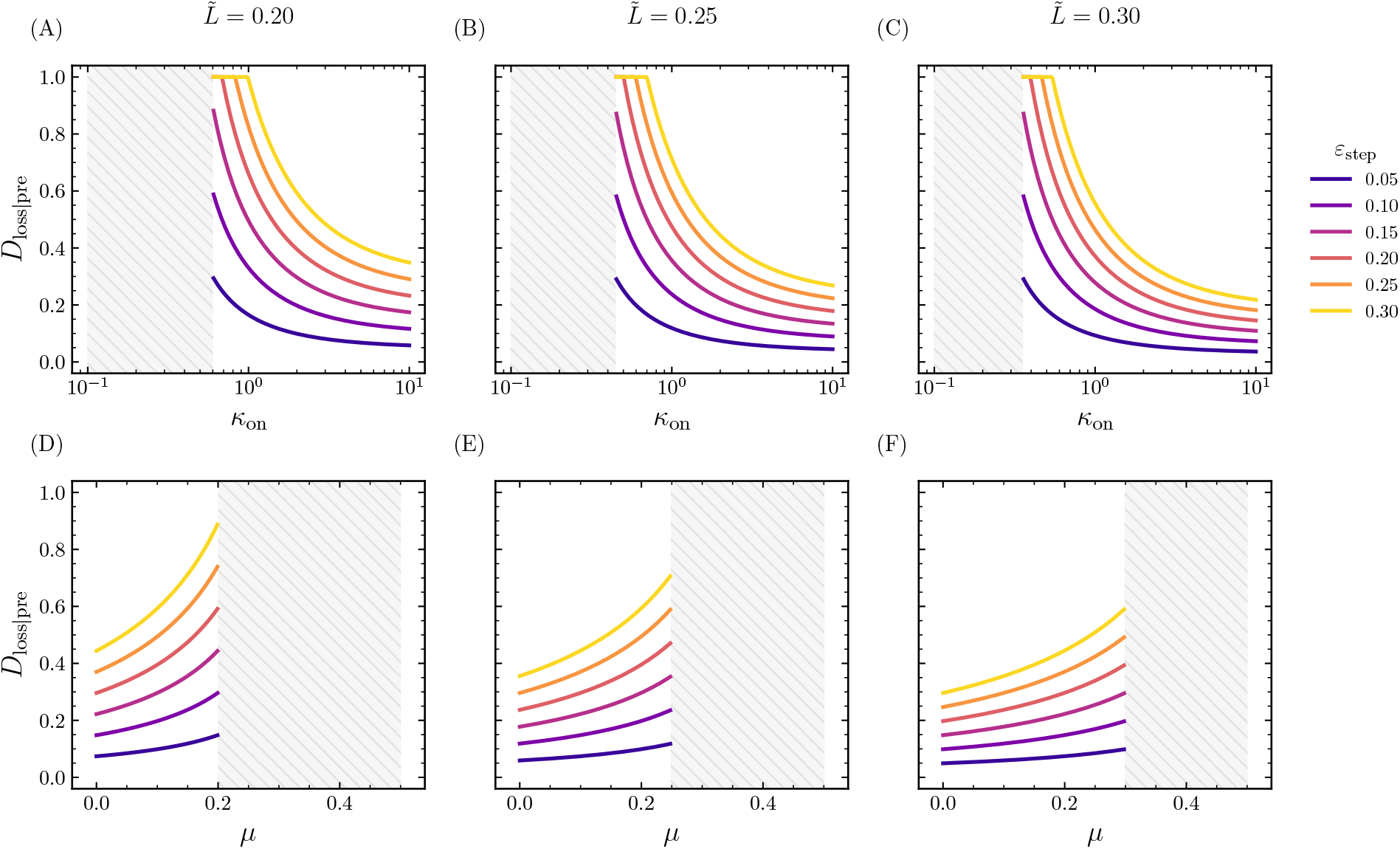
Post-stretch loss analysis of the reduced model across ACP binding and contractile shortening sweeps. Relative post-stretch loss *D*_loss|pre_ predicted by the reduced model for selected filament lengths 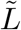 and step-strain amplitudes *ε*_step_. Panels show sweeps over *κ*_on_ **(A–C)** and *µ* **(D–F)** for 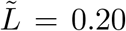, 0.25, and 0.30. Gray hatched regions indicate pre-stretch non-spanning conditions excluded from the loss analysis.

**Fig. S5.**
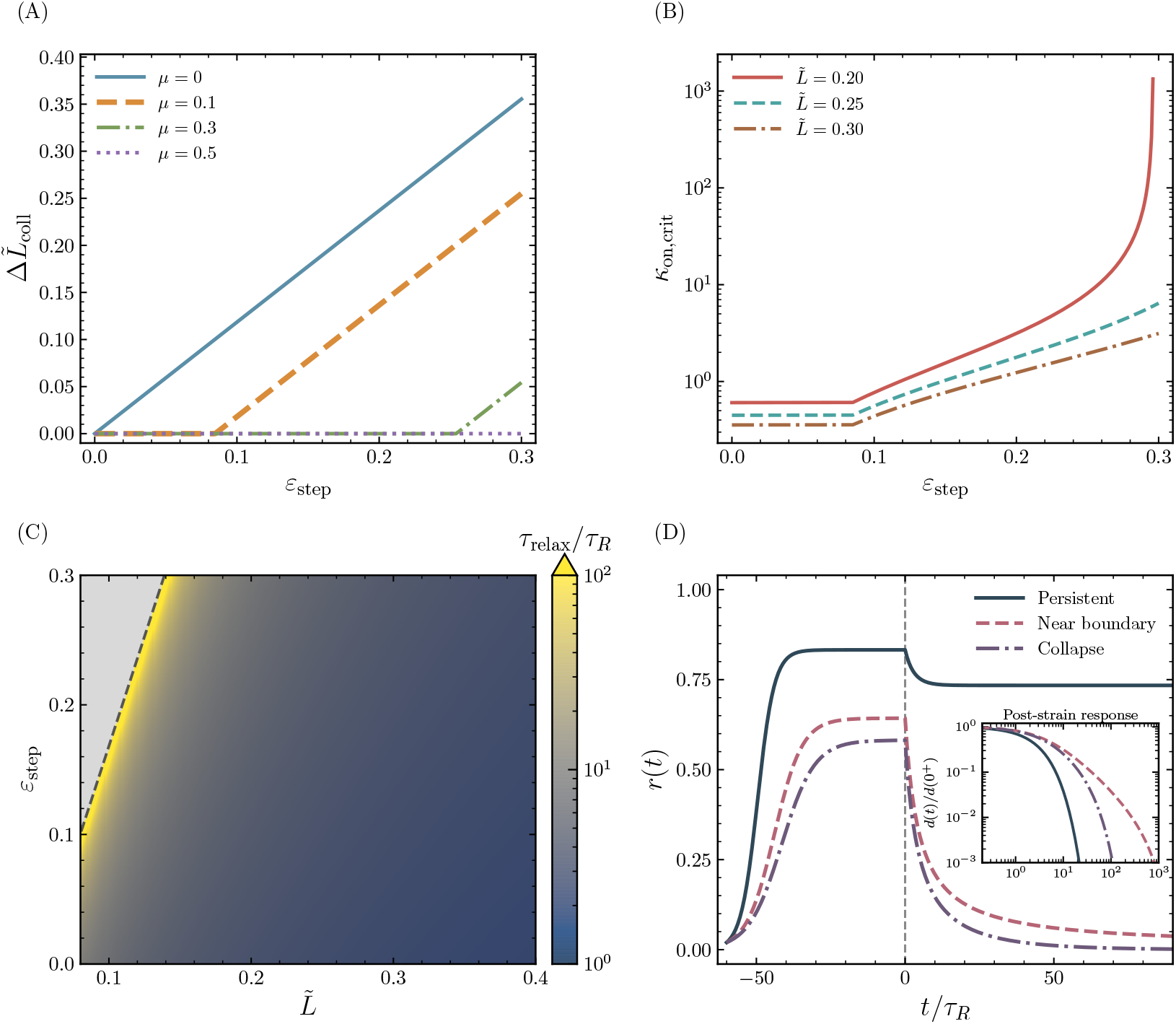
The reduced model links collapse-window width, rebinding rescue, and slow relaxation near the collapse–persistent boundary. **(A)** Collapse-window width 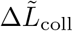 versus step-strain amplitude *ε*_step_ for different values of actomyosin-induced contractile shortening *µ*. **(B)** Critical ACP binding rate 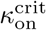 for maintaining persistence versus step-strain amplitude *ε*_step_ for different filament lengths 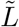. **(C)** Heatmap of the normalized linear relaxation timescale *τ*_relax_*/τ*_*R*_ in the (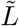, *ε*_step_) plane, computed by linearizing near the stable formed fixed point. The gray region indicates conditions without a stable formed fixed point, and the dashed line marks the corresponding collapse–persistent boundary. **(D)** Representative trajectories of the reduced connectivity variable *r*(*t*) before and after a step strain, illustrating persistent, near-boundary slow, and collapse responses. The gray dashed line marks the step strain input at *t* = 0. The inset shows the distance from the post-strain fixed point normalized by its value immediately after the strain step, 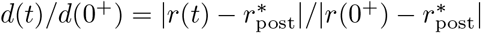, on a logarithmic scale.

